# Characterization of a Y-specific duplication/insertion of the anti-Mullerian hormone type II receptor gene based on a chromosome-scale genome assembly of yellow perch, *Perca flavescens*

**DOI:** 10.1101/717397

**Authors:** Romain Feron, Margot Zahm, Cédric Cabau, Christophe Klopp, Céline Roques, Olivier Bouchez, Camille Eché, Sophie Valière, Cécile Donnadieu, Pierrick Haffray, Anastasia Bestin, Romain Morvezen, Hervé Acloque, Peter T. Euclide, Ming Wen, Elodie Jouano, Manfred Schartl, John H. Postlethwait, Claire Schraidt, Mark R. Christie, Wes Larson, Amaury Herpin, Yann Guiguen

## Abstract

**Background:** Yellow perch, *Perca flavescens*, is an ecologically and commercially important species native to a large portion of the northern United States and southern Canada. It is also a promising candidate species for aquaculture. No yellow perch reference genome, however, has been available to facilitate improvements in both fisheries and aquaculture management practices.

**Findings:** By combining Oxford Nanopore Technologies long-reads, 10X genomics Illumina short linked reads and a chromosome contact map produced with Hi-C, we generated a high-continuity chromosome scale yellow perch genome assembly of 877.4 Mb. It contains, in agreement with the known diploid chromosome yellow perch count, 24 chromosome-size scaffolds covering 98.8% of the complete assembly (N50 = 37.4 Mb, L50 = 11). Genome annotation identified 41.7% (366 Mb) of repeated elements and 24,486 genes including 16,579 genes (76.3%) significantly matching with proteins in public databases. We also provide a first characterization of the yellow perch sex determination locus that contains a male-specific duplicate of the anti-Mullerian hormone type II receptor gene (*amhr2by*) inserted at the proximal end of the Y chromosome (chromosome 9). Using this sex-specific information, we developed a simple PCR genotyping test which accurately differentiates XY genetic males (*amhr2by*^+^) from XX genetic females (*amhr2by*^−^).

**Conclusions:** Our high-quality genome assembly is an important genomic resource for future studies on yellow perch ecology, toxicology, fisheries, and aquaculture research. In addition, the characterization of the *amhr2by* gene as a candidate sex determining gene in yellow perch provides a new example of the recurrent implication of the transforming growth factor beta pathway in fish sex determination, and highlights gene duplication as an important genomic mechanism for the emergence of new master sex determination genes.

## DATA DESCRIPTION

### Introduction and background

Yellow perch, *Perca flavescens* (Figure 1), is an ecologically and economically important species native to a large portion of the northern United States and southern Canada. Yellow perch supports recreational and commercial fisheries and is a major component of the food web in many inland lakes, where they are often the most abundant prey for larger species such as walleye (*Sander vitreus*), northern pike (*Esox lucius*), muskellunge (*Esox masquinongy*), and lake trout (*Salvelinus namaycush*) [1]. In the Laurentian Great Lakes, yellow perch are an important native species that has been heavily impacted by fishing pressure and environmental changes over the last century [2, 3]. Yellow perch is consistently among the most valuable commercially harvested fish species in the Great Lakes ($2.64/lb. dockside value in 2000 [4]), with fillets selling as high as $12/lb). However, many yellow perch fisheries have been forced to close due to substantial population declines [5]. The mechanisms underlying these declines are not fully understood but could be investigated using a combination of ecological and genetic studies if adequate genomic information were available.

**Figure 1:**
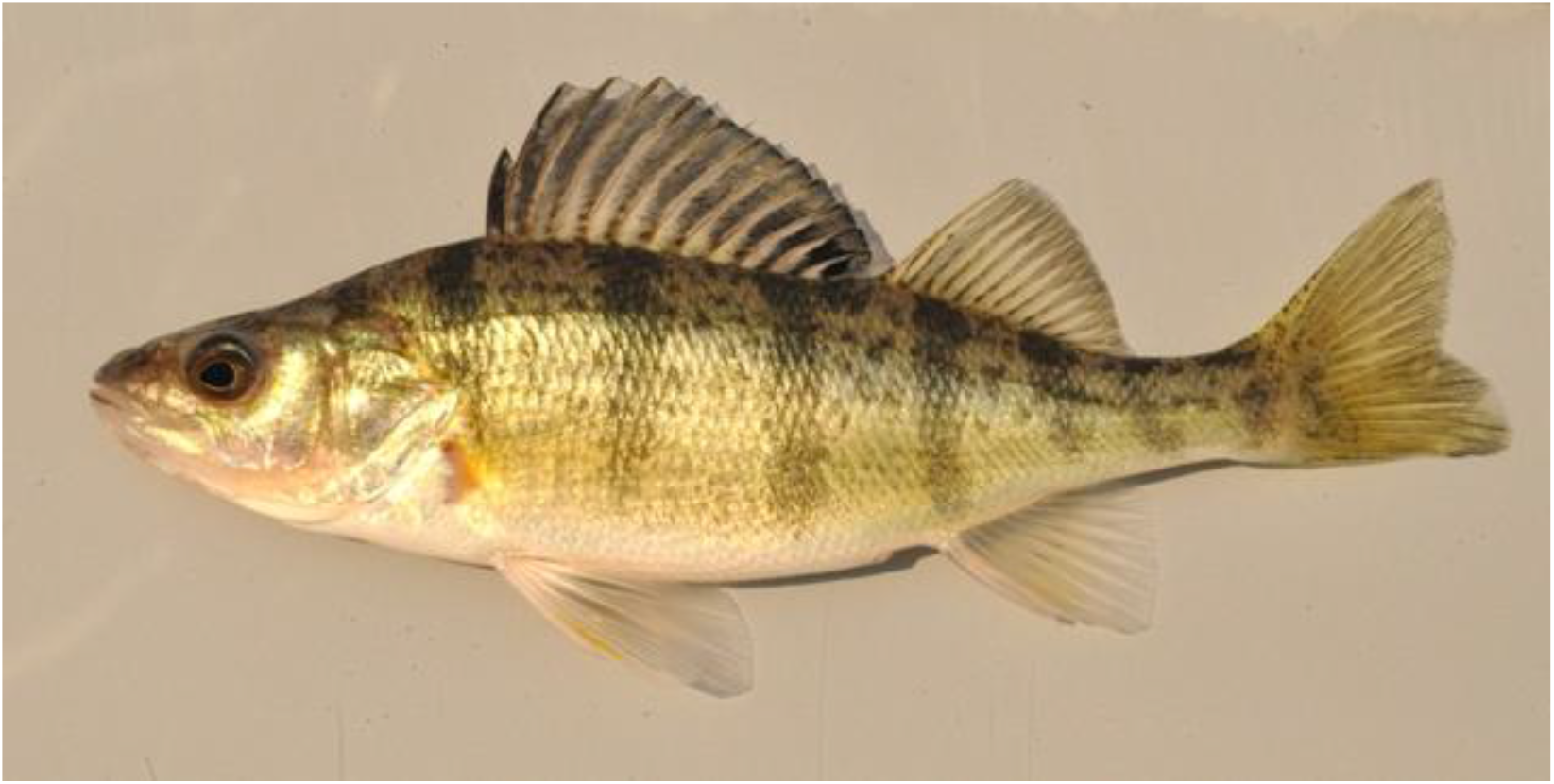
Adult yellow perch (*Perca flavescens*).

From an aquaculture perspective, yellow perch has many desirable attributes. For example, yellow perch can tolerate high stocking densities, are relatively disease resistant, and can be raised successfully under a variety of temperature and water conditions [6, 7]. Furthermore, yellow perch can be reared from hatching to marketable size in a relatively short period of time (∼1 year *vs*. 2+ years for most salmonids). Because yellow perch eat a diverse array of prey items [8], their feed can be obtained from ecologically sustainable sources while remaining cost effective (in contrast salmon are often fed a diet consisting primarily of other wild-caught fishes, known as fish meal). Lastly, yellow perch fillets have a firm texture and a mild flavor yielding a high market value.

The challenges faced by the yellow perch aquaculture include: increasing the spawning window for broodstock, reducing early life stage mortality, and developing large-bodied strains with faster growth rates [6]. Yellow perch spawn seasonally (typically in late spring to early summer) during a relatively narrow period of time (1-2 weeks). From an aquaculture perspective, it can be challenging to find males and females that are ready to spawn at the same time and, if the fish are not monitored daily, the peak spawning period can be missed entirely [9]. Compared to other aquaculture species, yellow perch also have a protracted free-swimming larval stage (∼30 days), during which the fish require precise food and water conditions for optimal survival. Developing broodstock that produce offspring with a shorter larval stage or that produce larger, more robust offspring would allow perch to be successfully reared in a broader array of facilities. Lastly, while yellow perch can already be grown to marketable size relatively quickly, the relative lack of selective breeding means that there is considerable room for developing yellow perch strains with faster growth rates and larger body sizes [6].

These challenges, which currently limit the wide-scale adoption of yellow perch as an aquaculture species, can be addressed using cutting edge genomic resources, such as the genome assembly described here. For example, one straightforward step towards obtaining fish with faster growth rates and larger body size would be to produce genetically all-female populations, as females grow considerably faster and larger than males [10–12]. More generally, sequencing and characterizing the yellow perch genome will facilitate improvements in both aquaculture and fisheries management practices.

### Results and Discussion

#### Genome characteristics

By a combination of three approaches -- Oxford Nanopore Technologies (ONT) long-reads, 10X genomics Illumina short linked reads (PE150 chemistry), and a chromosome contact map (Hi-C) -- we generated a high-continuity, chromosome length *de novo* genome assembly of the yellow perch. Before the Hi-C integration step, the assembly yielded a genome size of 877 Mb with 879 contigs, a N50 contig size of 4.3 Mb, and a L50 contig number of 60 (i.e. half of the assembled genome is included in the 60 longest contigs). After Hi-C integration, the genome assembled into 269 fragments with a total length of 877.4 Mb, including 24 chromosome-length scaffolds representing 98.78 % of the complete genome sequence (N50 = 37.4 Mb, L50 = 11) (see Table 1). Genome sizes are both very close to the 873 Mbp GenomeScope [13] estimation based on short-read analysis with a repeat length of 266 Mbp (30.5%) and slightly lower than the estimation of *P. flavescens* genome sizes based on C-values (900 Mbp and 1200 Mbp records in the Animal Genome Size Database [14]). The 24 chromosome-length scaffolds obtained after Hi-C integration are consistent with the diploid chromosome (Chr) number of yellow perch (2n = 48) [15]. The genome completeness of these assemblies was estimated using Benchmarking Universal Single-Copy Orthologs (BUSCO) v3.0 [16] based on the Actinopterygii database. BUSCO scores (see Table 1) of the pre-Hi-C and post-Hi-C assemblies were roughly similar (Complete BUSCOs between 97.6% and 97.8%) and with small values for both fragmented (< 1%) and missing (< 1.5%) BUSCO genes.

**Table 1.**
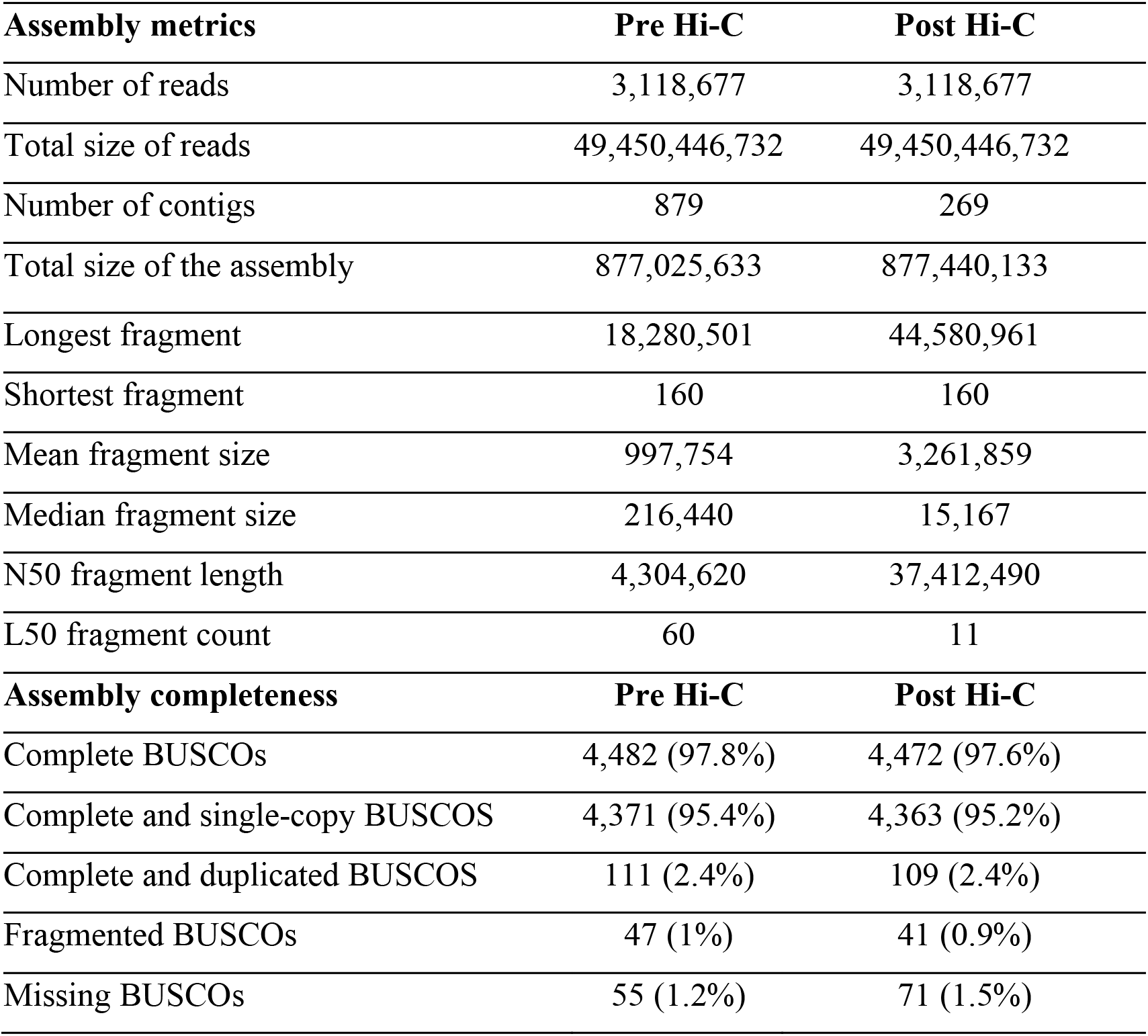
Yellow perch assembly statistics.

| Assembly metrics | Pre Hi-C | Post Hi-C |
| --- | --- | --- |
| Number of reads | 3,118,677 | 3,118,677 |
| Total size of reads | 49,450,446,732 | 49,450,446,732 |
| Number of contigs | 879 | 269 |
| Total size of the assembly | 877,025,633 | 877,440,133 |
| Longest fragment | 18,280,501 | 44,580,961 |
| Shortest fragment | 160 | 160 |
| Mean fragment size | 997,754 | 3,261,859 |
| Median fragment size | 216,440 | 15,167 |
| N50 fragment length | 4,304,620 | 37,412,490 |
| L50 fragment count | 60 | 11 |
| Assembly completeness | Pre Hi-C | Post Hi-C |
| Complete BUSCOs | 4,482 (97.8%) | 4,472 (97.6%) |
| Complete and single-copy BUSCOS | 4,371 (95.4%) | 4,363 (95.2%) |
| Complete and duplicated BUSCOS | 111 (2.4%) | 109 (2.4%) |
| Fragmented BUSCOs | 47 (1%) | 41 (0.9%) |
| Missing BUSCOs | 55 (1.2%) | 71 (1.5%) |

Repeated elements accounted for 41.71% (366 Mbp) of our chromosomal assembly and these regions were soft masked before gene annotation. Using protein, transcript, and *de novo* gene prediction evidence we annotated 24,486 genes, including 16,579 (76.3%) that significantly matched with a protein hit in the non-redundant NCBI database (Table 2). Our yellow perch genome was also annotated with the NCBI Eukaryotic Genome Annotation Pipeline (NCBI *Perca flavescens* Annotation Release 100 [17]), leading to a higher gene count (28,144) with possibly multiple transcripts per gene (Table 2).

**Table 2.**
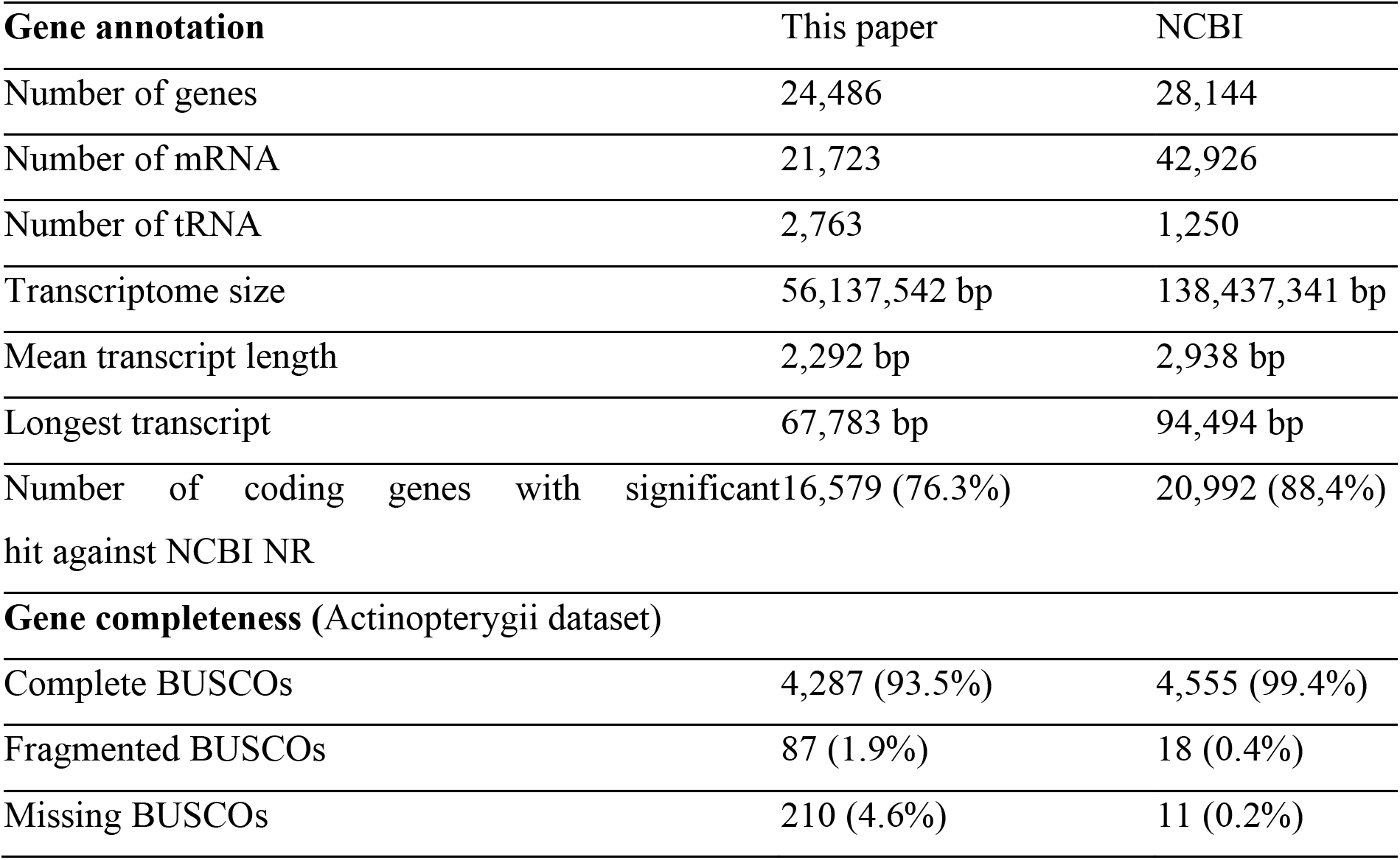
Yellow perch annotation statistics.

| **Gene annotation** | This paper | NCBI |
| --- | --- | --- |
| Number of genes | 24,486 | 28,144 |
| Number of mRNA | 21,723 | 42,926 |
| Number of tRNA | 2,763 | 1,250 |
| Transcriptome size | 56,137,542 bp | 138,437,341 bp |
| Mean transcript length | 2,292 bp | 2,938 bp |
| Longest transcript | 67,783 bp | 94,494 bp |
| Number of coding genes with significant hit against NCBI NR | 16,579 (76.3%) | 20,992 (88.4%) |
| **Gene completeness (Actinopterygii dataset)** |  |  |
| Complete BUSCOs | 4,287 (93.5%) | 4,555 (99.4%) |
| Fragmented BUSCOs | 87 (1.9%) | 18 (0.4%) |
| Missing BUSCOs | 210 (4.6%) | 11 (0.2%) |

#### Yellow perch sex-determination

Yellow perch has a male monofactorial heterogametic sex determination system (XX/XY) [18] with undifferentiated sex chromosomes [19]. Using a male-versus-female pooled gDNA whole genome sequencing strategy [20], we identified a relatively small region of 100 kb localized at the proximal end of chromosome 9 (Chr09:0-100,000 bp) with a complete absence of female reads, excluding repeated elements (Fig. 2.A-B). This coverage bias strongly supports the hypothesis that Chr09 is the yellow perch sex chromosome and contains a small Y-specific region in phenotypic males that is completely absent from phenotypic females. Genome annotation shows that this Y-specific insertion on Chr09 contains a duplicate copy (*amhr2by*) of the autosomal anti-Mullerian hormone receptor gene located on Chr04 (*amhr2a*). The *amhr2* gene has previously been characterized as a master sex-determining gene in some pufferfishes [21, 22] and the *hotei* mutation in the medaka *amhr2* gene induces a male-to-female sex-reversal of genetically XY fish [23]. However, in contrast to pufferfishes, in which the differentiation of X and Y chromosomes is extremely limited and originated from an allelic diversification process, the yellow perch *amhr2by* sequence is quite divergent from its *amhr2a* autosomal counterpart. Specifically, the *amhr2by* gene shows only 88.3 % identity with *amhr2a* in the aligned coding sequence and 89.1 % in the aligned parts of the introns, but with many long gaps and indels in the introns (Fig. 2C-D). This nucleotide sequence divergence impacts the protein sequence of the yellow perch *amhr2by* gene (Fig. 2D-2E), but due to a complete absence of exons 1 & 2 (Fig. 2C-2E) compared to its autosomal counterpart, the yellow perch Amhr2by protein translates as a N-terminal-truncated type II receptor that lack most of the cysteine-rich extracellular part of the receptor, which is crucially involved in ligand binding specificity [24].

**Figure 2:**
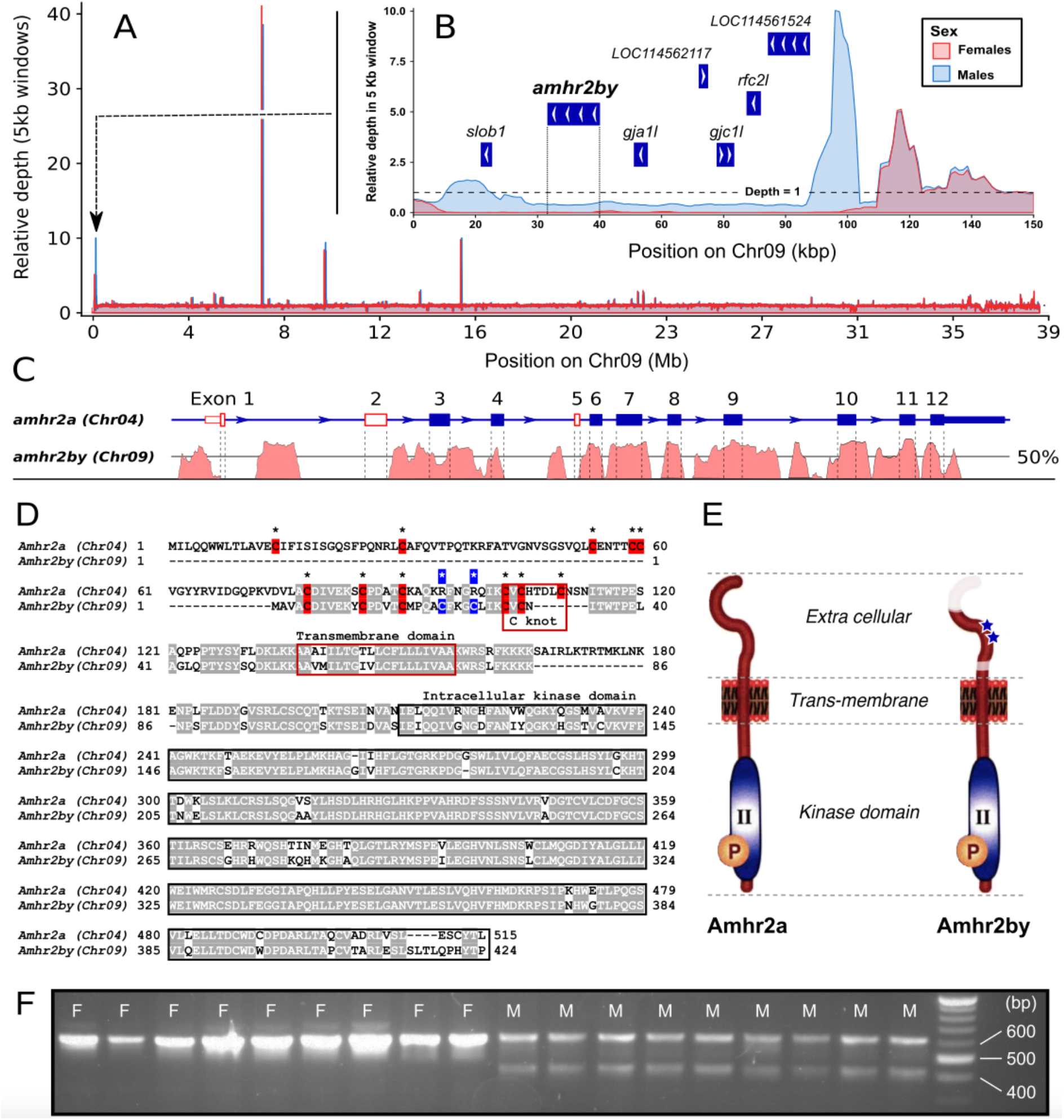
Characterization of a Y-specific duplication/insertion of the anti-Mullerian hormone receptor gene (*amhr2by*) in yellow perch. **A**. Pool-seq data illustrating relative read depth across chromosome 9 (Chr09) for the male (blue line) and female (red line) pools showing a coverage difference between males (blue area) and females (red area) in the first 100 kb of Chr09. **B.** Zoom-in on the read depth difference between males and females in the first 150 kb of Chr09. Gene annotation is represented by blue boxes with arrows to indicate transcript orientation (NCBI *Perca flavescens* Annotation Release 100 [13]). Abbreviations: *slob1* (probable inactive serine/threonine-protein kinase slob1, *LOC114561790*), *amhr2by* (anti-Mullerian hormone type-2 receptor-like, *LOC114561927*), *gja1l* (gap junction alpha-1 protein-like, *LOC114562210*), *gjcl1* (gap junction gamma-1 protein-like, *LOC114562012*), *rfc2l* (replication factor C subunit 2-like, *LOC114561955*). **C.** Identity plot of the alignment of *amhr2by* gene sequence (Chr09, bottom panel) with the autosomal *amhr2a* gene sequence (Chr04, top panel). The structure of the *amhr2a* gene is depicted with blue boxes (exons, E1 to E12) and blue lines (introns) with arrows indicating the transcription orientation. The solid line on the identity plot (bottom panel) represents 50% nucleotide identity between the two sequences. **D.** ClustalW [62] alignment of Amhr2a and Amhr2by proteins. Identical amino-acids are shaded and the cysteines in the extracellular domain of Amhr2 are shown with bolded black asterisks. Additional cysteines specific to Amhr2by are highlighted in blue. The different domains of the receptor are boxed. **E.** Schematic representation of the two yellow perch Amhr2 proteins showing that the main differences impact the extracellular domain with parts missing in Amhr2by represented in white and the two additional cysteines represented by blue asterisks. **F.** Validation of *amhr2by* sex linkage in yellow perch. Agarose gel electrophoresis of multiplex PCR of *amhr2a* (higher size PCR fragment, 638 bp), and *amhr2by* (lower size PCR fragment, 443 bp) in nine females (F, left side) and nine males (M, right side) genomic DNA.

To validate the male specificity of this potential Y-specific insertion, we designed primers specific for both *amhr2by* and *amhr2a* and genotyped 25 male and 25 female yellow perch collected from a Southeastern Lake Michigan population, which is geographically isolated from the Plum Lake (Wisconsin) population of the 30 males and 30 females used for initial analysis with pool-sequencing. The presence/absence of the *amhr2by* PCR product was perfectly correlated with the determined phenotypic sex, with the amplification of an *amhr2by* fragment only in the 25 males and no amplification in the 25 females (see Fig. 2F for 18 of the 50 individuals tested). The simultaneous amplification of the *amhr2a* fragment in both males and in females provided an internal control preventing single-locus dropout in such a multiplexed PCR reaction.

This complete sex-linkage result makes the yellow perch *amhr2by* an obvious candidate as a sex determining gene. Interestingly, anti-Mullerian hormone (Amh) has been also characterized as a male-promoting gene in zebrafish [25] and as a master sex determining gene both in Patagonian pejerrey [26], Nile tilapia [27] and Northern pike [28]. The role of transforming growth factor beta (TGF-ß) members in sex determination is not limited to the Amh pathway; additional TGF-ß family genes have also been characterized as master sex determining genes, including growth differentiation factor 6 (*gdf6*) in the turquoise killifish [29] and gonadal soma derived factor (*gsdf*) in the Luzon medaka and the sablefish [30, 31]. Additional evidence, including loss of *amhr2by* function experiments in XY males and gain of *amhr2by* function experiments in XX females, is necessary to critically test the hypothesis that this male-specific *amhr2by* duplication really functions as a master sex determining gene in yellow perch. However, given the known importance of the Amh pathway in fish sex determination, and that no other gene in that small sex locus is known to play a role in sex differentiation, *amhr2by* is a prime candidate for the yellow perch master sex determining gene. This finding provides another example of the recurrent utilization of the TGF-ß pathway in fish sex determination, and thus supports the ‘limited option’ hypothesis [32], which states that some genes are more likely than others to be selected as master sex determining genes. How this N terminal truncated Amhr2 could trigger its function as a master sex determining gene is as yet unknown, but one hypothesis is that this truncation constitutively activates the Amh receptor causing it to signal in the absence of Amh ligand.

However, regardless of the precise role of the structural variation of amhr2 in sex determination, we have developed a simple molecular protocol for genotypically sexing perch of any life stage and produced a fully annotated, chromosome-scale genome assembly that will undoubtedly aid in the conservation and management of this species.

## MATERIAL AND METHODS

### Sampling and genomic DNA extraction

The male yellow perch used for whole genome sequencing was sampled from Plum Lake, Vilas County, Wisconsin, USA. A 0.5 ml blood sample was taken from this animal and immediately put in a TNES-Urea lysis buffer (TNES-Urea: 4 M urea; 10 mM Tris-HCl, pH 7.5; 125 mM NaCl; 10 mM EDTA; 1% SDS) [33]. High molecular weight genomic DNA (gDNA) was then purified by phenol-chloroform extraction. For the chromosome contact map (Hi-C), 1.5 ml of blood was taken from a different male from a domesticated line of yellow perch raised at the Farmory, an aquaculture facility in Green Bay, Wisconsin, USA. The fresh blood sample was slowly cryopreserved with 15 % Dimethyl sulfoxide (DMSO) in a Mr. Frosty Freezing Container (Thermo Scientific) at −80°C. Fin clip samples (30 males and 30 females) for whole-genome sequencing of pools of individuals (Pool-seq) were collected from wild yellow perch in Green Bay, Lake Michigan, Wisconsin, USA, placed in 90% ethanol and then stored dried until gDNA extraction was performed using the NucleoSpin Kit for Tissue (Macherey-Nagel, Duren, Germany). Genomic DNAs from individual fish were then quantified using a Qubit fluorometer (Thermofisher) and pooled in equimolar ratios by individual and sex, resulting in one gDNA pool for males and one gDNA pool for females. For validation of *amhr2by* sex-linkage, 50 phenotypically sexed individuals (25 males and 25 females) wild perch from Lake Michigan were collected in May of 2018 using gill net sets off the shore of Michigan City, Indiana (41°42.5300’N, 86°57.5843’W). Upon collection, each individual fish was euthanized, phenotypic sex was determined by visual inspection of gonads during necropsy, and caudal fin clips were taken from each yellow perch individual and stored in 95% non-denatured ethanol. Genomic DNA was extracted using the DNeasy extraction kit and protocol (Qiagen).

### DNA library construction and sequencing

#### Nanopore sequencing

The quality and purity of gDNA was assessed using spectrophotometry, fluorometry and capillary electrophoresis. Additional purification steps were performed using AMPure XP beads (Beckman Coulter). All library preparations and sequencing were performed using Oxford Nanopore Ligation Sequencing Kits SQK-LSK108 (Oxford Nanopore Technology) (14 flowcells) or SQK-LSK109 (2 flowcells) according to the manufacturer’s instructions. For the SQK-LSK108 sequencing Kit, 140 μg of DNA was purified and then sheared to 20 kb using the megaruptor system (Diagenode). For each library, a DNA-damage-repair step was performed on 5 μg of DNA. Then an END-repair+dA-tail-of-double-stranded-DNA-fragments step was performed and adapters were ligated to DNAs in the library. Libraries were loaded onto two R9.5 and twelve R9.4 flowcells and sequenced on a GridION instrument at a concentration of 0.1 pmol for 48 h. For the SQK-LSK109 sequencing Kit, 10 μg of DNA was purified and then sheared to 20 kb using the megaruptor system (Diagenode). For each library, a one-step-DNA-damage repair+END-repair+dA-tail-of-double-stranded-DNA-fragments procedure was performed on 2 μg of DNA. Adapters were then ligated to DNAs in the library. Libraries were loaded on R9.4.1 flowcells and sequenced on either a GridION or PromethION instrument at a concentration of 0.05 pmol for 48h or 64h respectively. The 15 GridION flowcells produced 69.4 Gb of data and the PromethION flowcell produced 65.5 Gb of data.

#### 10X Genomics sequencing

The Chromium library was prepared according to 10X Genomics’ protocols using the Genome Reagent Kit v2. Sample quantity and quality controls were validated by Qubit, Nanodrop and Femto Pulse machines. The library was prepared from 10 ng of high molecular weight (HMW) gDNA. Briefly, in the microfluidic Genome Chip, a library of Genome Gel Beads was combined with HMW template gDNA in master mix and partitioning oil to create Gel Bead-In-EMulsions (GEMs) in the Chromium apparatus. Each Gel Bead was then functionalized with millions of copies of a 10x™ barcoded primer. Dissolution of the Genome Gel Bead in the GEM released primers containing (i) an Illumina R1 sequence (Read 1 sequencing primer), (ii) a 16 bp 10x Barcode, and (iii) a 6 bp random primer sequence. R1 sequence and the 10x™ barcode were added to the molecules during the GEM incubation. P5 and P7 primers, R2 sequence, and Sample Index were added during library construction. 10 cycles of PCR were applied to amplify the library. Library quality was assessed using a Fragment Analyser and library was quantified by qPCR using the Kapa Library Quantification Kit. The library was then sequenced on an Illumina HiSeq3000 using a paired-end read length of 2×150 nt with the Illumina HiSeq3000 sequencing kits and produced 315 million read pairs.

#### Hi-C sequencing

*In situ* Hi-C was performed according to previously described protocols [34]. Cryopreserved blood cells were defrosted, washed with PBS twice and counted. 5 million cells were then cross-linked with 1% formaldehyde in PBS, quenched with Glycine 0.125M and washed twice with PBS. Membranes were then disrupted with a Dounce pestle, nuclei were permeabilized using 0.5% SDS and then digested with *Hind*III endonuclease. 5’-overhangs at *Hind*III-cut restriction sites were filled-in, in the presence of biotin-dCTP with the Klenow large fragment, and then re-ligated at a NheI restriction site. Nuclei were lysed and DNA was precipitated and then purified using Agencourt AMPure XP beads (Beckman Coulter) and quantified using the Qubit fluorometric quantification system (Thermo). T4 DNA polymerase was used to remove un-ligated biotinylated ends. Then the Hi-C library was prepared according to Illumina’s protocols using the Illumina TruSeq Nano DNA HT Library Prep Kit with a few modifications: 1.4μg DNA was fragmented to 550nt by sonication. Sheared DNA was then sized (200-600pb) using Agencourt AMPure XP beads, and biotinylated ligation junctions were captured using M280 Streptavidin Dynabeads (Thermo) and then purified using reagents from the Nextera Mate Pair Sample preparation kit (Illumina). Using the TruSeq nano DNA kit (Illumina), the 3’ ends of blunt fragments were adenylated. Next, adaptors and indexes were ligated and the library was amplified for 10 cycles. Library quality was assessed by quantifying the proportion of DNA cut by endonuclease NheI using a Fragment Analyzer (Advanced Analytical Technologies, Inc., Iowa, USA). Finally, the library was quantified by qPCR using the Kapa Library Quantification Kit (Roche). Sequencing was performed on an Illumina HiSeq3000 apparatus (Illumina, California, USA) using paired-end 2×150 nt reads. This produced 128 million read pairs (38.4 Gb of raw nucleotides).

#### Pool sequencing

Pool-sequencing libraries were prepared according to Illumina’s protocols using the Illumina TruSeq Nano DNA HT Library Prep Kit (Illumina, California, USA). In short, 200 ng of each gDNA pool (males and females pools) was fragmented to 550 bp by sonication on M220 Focused-ultrasonicator (COVARIS). Size selection was performed using SPB beads (kit beads) and the 3’ ends of blunt fragments were mono-adenylated. Then, adaptors and indexes were ligated and the construction was amplified with Illumina-specific primers. Library quality was assessed using a Fragment Analyzer (Advanced Analytical Technologies, Inc., Iowa, USA) and libraries were quantified by qPCR using the Kapa Library Quantification Kit (Roche). Sequencing was performed on a NovaSeq (Illumina, California, USA) using a paired-end read length of 2×150 nt with Illumina NovaSeq Reagent Kits. Sequencing produced 119 million paired reads for the male pool library and 132 million paired reads for the female pool library.

### Genome assembly and analysis

#### Genome size estimation

K-mer-based estimation of the genome size was carried out with GenomeScope [13]. 10X reads were processed with Jellyfish v1.1.11 [35] to count 17-, 19-, 21-, 23- and 25-mers with a max k-mer coverage of 10,000.

#### Genome assembly

GridION and PromethION data were trimmed using Porechop v0.2.1 [36], corrected using Canu v1.6 [37] and filtered to keep only reads longer than 10 kbp. Corrected reads were then assembled using SmartDeNovo version of May-2017 [38] with default parameters. The assembly base pair quality was improved by several polishing steps including two rounds of long read alignment to the draft genome with minimap2 v2.7 [39] followed by Racon v1.3.1 [40], as well as three rounds of 10X genomics short read alignments using Long Ranger v2.1.1 (10x Genomics 2018) followed by Pilon v1.22 [41]. The polished genome assembly was then scaffolded using Hi-C as a source of linking information. Reads were aligned to the draft genome using Juicer [42] with default parameters. A candidate assembly was then generated with 3D de novo assembly (3D-DNA) pipeline [43] with the -r 0 parameter. Finally, the candidate assembly was manually reviewed using Juicebox Assembly Tools [42]. Genome completeness was estimated using Benchmarking Universal Single-Copy Orthologs (BUSCO) v3.0 [16] based on 4,584 BUSCO orthologs derived from the Actinopterygii lineage.

#### Genome annotation

The first annotation step was to identify repetitive content using RepeatMasker v4.0.7 [43], Dust (Kuzio et al., unpublished but described in [44]), and TRF v4.09 [45]. A species-specific *de novo* repeat library was built with RepeatModeler v1.0.11 [46] and repeated regions were located using RepeatMasker with the *de novo* and *Danio rerio* libraries. Bedtools v2.26.0 [47] was used to merge repeated regions identified with the three tools and to soft mask the genome. The MAKER3 genome annotation pipeline v3.01.02-beta [48] combined annotations and evidence from three approaches: similarity with fish proteins, assembled transcripts (see below), and *de novo* gene predictions. Protein sequences from 11 fish species (*Astyanax mexicanus*, *Danio rerio*, *Gadus morhua*, *Gasterosteus aculeatus*, *Lepisosteus oculatus*, *Oreochromis niloticus*, *Oryzias latipes*, *Poecilia formosa*, *Takifugu rubripes*, *Tetraodon nigroviridis*, *Xiphophorus maculatus*) found in Ensembl were aligned to the masked genome using Exonerate v2.4 [49]. As *Perca fluviatilis* is a relatively closely related species from *P. flavescens* (divergence time is estimated to be 19.8 million years ago according to [50]), RNA-Seq reads of *P. fluviatilis* (NCBI BioProject PRJNA256973) from the PhyloFish project [51] were used for genome annotation and aligned to the chromosomal assembly using STAR v2.5.1b [52] with outWigType and outWigStrand options to output signal wiggle files. Cufflinks v2.2.1 [53] was used to assemble the transcripts which were used as RNA-seq evidence. Braker v2.0.4 [54] provided *de novo* gene models with wiggle files provided by STAR as hint files for GeneMark and Augustus training. The best supported transcript for each gene was chosen using the quality metric called Annotation Edit Distance (AED) [55]. Genome annotation gene completeness was assessed by BUSCO using the Actinopterygii group. Finally, predicted genes were subjected to similarity searches against the NCBI NR database using Diamond v0.9.22 [56]. The top hit with a coverage over 70% and identity over 80% was retained.

#### Pool-sequencing analysis

Reads from the male and female pools were aligned to the chromosomal assembly with BWA mem (version 0.7.17, [57]), and the resulting BAM files were sorted and PCR duplicates removed using Picard tools (version 2.18.2). A file containing the nucleotide composition of each pool for each genomic position was generated using samtools mpileup (version 1.8, [58]) and popoolation2 mpileup2sync (version 1201, [59]). This file was then analyzed with custom software (PSASS version 2.0.0 [60]) to compute 1) the position and density of sex-specific SNPs, defined as SNPs heterozygous in one sex while homozygous in the other sex, 2) absolute and relative read depth for the male and female pools along the genome, and 3) F_ST_ between males and females in windows along the genome. PSASS was run with default parameters except --window-size which was set to 5000 and --output-resolution which was set to 1000.

#### Validation of *amhr2by* sex-linkage

To validate the sex-linkage of *amhr2by* in males suggested by the pool-sequencing results, two primer sets were designed based on the alignment of yellow perch *amhr2a* and *amhr2by* genes with one primer pair specific for the autosomal *amhr2a* gene (forward: 5’-GGGAAACGTGGGAAACTCAC-3’, and reverse: 5’-AGCAGTAGTTACAGGGCACA-3’, expected fragment size: 638 bp) and one primer pair specific for the Y chromosomal *amhr2by* gene (forward: 5’-TGGTGTGTGGCAGTGATACT-3’, and reverse: 5’-ACTGTAGTTAGCGGGCACAT-3’, expected fragment size: 443 bp). Gene alignments were run with mVISTA [61]. Primers were sourced from Integrated Data Technologies (IDT). All samples were run blind with respect to phenotypic sex; the male and female samples were randomized, and their phenotypic sex was not cross referenced with field data until gel electrophoresis was run on the final PCR products. Genotyping was carried out on each gDNA sample using a multiplexed PCR approach. The PCR reaction solution was composed of 50 μl of PCR Master Mix (Quiagen), 10 μl of each primer (40 μl total), and 10 μl of gDNA (concentrations of gDNA ranging from 150 to 200 ng/ μl) for a total reaction volume of 100 μl. Thermocycling conditions were 1 cycle of 3 min at 94°C, followed by 35 cycles of 30 sec at 94°C, 30 sec at 51°C, and 1 min at 72°C, and finishing with 10 min incubation at 72°C. PCR products were loaded on a 1.5 % agarose gel, run at 100V for 45 minutes and visualized with a UVP UVsolo touch UV box.

## Availability of supporting data

This Whole Genome Shotgun project has been deposited at DDBJ/ENA/GenBank under the accession SCKG00000000. The version described in this paper is version SCKG01000000. Hi-C, 10X genomics and pool-sequencing Illumina reads, and Oxford Nanopore Technologies genome raw reads are available in the Sequence Read Archive (SRA), under BioProject reference PRJNA514308.

## Author contributions

YG, MS, and JHP designed the project. WL, CS and MC collected the samples, EJ, MW, CR, OB, SV and HA extract the gDNA, made the genomic libraries and sequenced them. RF, CC, CK, MZ, PE, AH and YG processed the genome assemblies and / or analyzed the results. CS and MC checked sex-linkage of *amhr2by* on yellow perch samples. YG, RF, WL, MC, JHP, CC, CK, and CR wrote the manuscript. MS, JHP, CD, PH, AB, RM, MC and YG, supervised the project administration and raised funding. All the authors read and approved the final manuscript.

## Acknowledgements

This project was supported by funds from the “Agence Nationale de la Recherche” and the “Deutsche Forschungsgemeinschaft” (ANR/DFG, PhyloSex project, 2014-2016), the CRB-Anim “Centre de Ressources Biologiques pour les animaux domestiques” project PERCH’SEX, the FEAMP “Fonds européen pour les affaires maritimes et la pêche” project SEX’NPERCH, and R01 GM085318 from the National Institutes of Health, USA. Additional funding was provided to MRC from the Great Lakes Fishery Commission, Project ID: 2018_CHR_44072. The GeT core facility, Toulouse, France was supported by France Génomique National infrastructure, funded as part of “Investissement d’avenir” program managed by Agence Nationale pour la Recherche (contract ANR-10-INBS-09). We are grateful to the Genotoul bioinformatics platform Toulouse Midi-Pyrenees (Bioinfo Genotoul) for providing computing and/or storage resources. Any use of trade, product, or company name is for descriptive purposes only and does not imply endorsement by the U.S. Government.

